# Role of HNF4alpha-cMyc Interaction in CDE-diet Induced Liver Injury and Regeneration

**DOI:** 10.1101/2023.11.27.568898

**Authors:** Manasi Kotulkar, Julia Barbee, Diego Paine Cabrera, Dakota Robarts, Udayan Apte

**Affiliations:** Department of Pharmacology, Toxicology, and Therapeutics, University of Kansas Medical Center, Kansas City, KS

**Keywords:** Liver regeneration, CDE diet, HNF4α

## Abstract

**Background:** Hepatocyte nuclear factor 4 alpha (HNF4α) is a nuclear factor essential for liver function and regeneration. HNF4α negatively regulates the expression of cMyc, which plays an important role in proliferation and differentiation during liver regeneration. This study investigated the role of HNF4α-cMyc interaction in regulating liver injury and regeneration using the choline-deficient and ethionine-supplemented (0.15%) (CDE) diet feeding model, which exhibits characteristics of chronic liver diseases including liver injury, inflammation, early fibrotic changes along with hepatocyte and biliary epithelial cell regeneration, and activation of hepatic progenitor cells (HPC).

**Methods:** Wild-type (WT), hepatocyte-specific knockout of HNF4α (HNF4α-KO), cMyc (cMyc-KO), and HNF4α-cMyc double knockout (DKO) mice were fed a CDE diet for one week to induce subacute liver injury. To study regeneration and recovery, mice were fed a one-week CDE diet followed by a one-week recovery period on a normal chow diet.

**Results:** WT mice showed significant liver injury and decreased HNF4α mRNA and protein expression after one week of a CDE diet. WT mice also showed an increase in markers of proliferation and HPC activation, but no major change in markers of inflammation or fibrosis.

The HNF4α-KO mice exhibited baseline hepatomegaly, which significantly declined during the recovery period. HNF4α deletion resulted in significantly higher injury compared to WT mice after one week of CDE diet feeding but similar recovery. Markers of inflammation, fibrosis, proliferation, and HPC activation were significantly higher in HNF4α-KO mice during the injury period but declined during the recovery period.

The cMyc-KO mice showed increased injury after one week of the CDE diet, but it was substantially lower than the WT and HNF4α-KO mice. Deletion of cMyc resulted in a significant activation of inflammatory genes higher than in the WT and HNF4α-KO mice. Whereas fibrosis and proliferation markers increased in cMyc-KO mice, they were substantially lower than in HNF4α-KO mice and similar to WT mice. cMyc-KO also showed an increase in HPC markers following one week of CDE-induced injury.

Deletion of both HNF4α and cMyc in DKO mice resulted in significant liver injury comparable to the HNF4α-KO mice after one week of CDE diet feeding, but led to complete recovery. Markers of inflammation, fibrosis, and proliferation increased after CDE diet feeding, were higher than WT mice, and comparable to HNF4α-KO mice. Interestingly, DKO mice showed a significant increase in HPC markers both following one week of CDE-induced injury and after one week of recovery.

**Conclusions:** These data indicate that deletion of HNF4α increases and deletion of cMyc decreases subacute liver injury induced by a one week CDE diet feeding. Deletion of HNF4α results in increased inflammation, fibrosis, proliferation, and HPC activation, all of which except inflammation are reduced following cMyc deletion. Simultaneous deletion of HNF4α and cMyc results in a phenotype similar to HNF4α deletion but with higher HPC activation. Taken together, these data show that HNF4α protects against inflammatory and fibrotic change following CDE diet-induced injury, which is driven by cMyc.

## Introduction

Hepatocyte nuclear factor 4 alpha (HNF4α) is a key nuclear receptor expressed by hepatocytes. HNF4α is important for liver development and the maintenance of mature liver function. HNF4α plays an important role in maintaining hepatocyte differentiation [1]. Decreased HNF4α results in the loss of hepatic function and a de-differentiated phenotype [2]. Ablation of HNF4α in the liver induces spontaneous proliferation of hepatocytes via induction of mRNA and protein expression of several pro-mitogenic genes, including cMyc [3]. A recent study from our lab investigated the role of HNF4α-cMyc interaction in liver regeneration after drug-induced acute liver injury. We found that HNF4α interacts with Nrf2 and contributes to recovery from acetaminophen-induced liver injury by downregulating the expression of cMyc [4].

The choline-deficient, ethionine-supplemented (CDE) diet is a model used to induce chronic liver injury in mice. Dietary deficiency of choline results in disrupted fat metabolism and the secretion of very low-density lipoproteins, contributing to steatosis. Ethionine is a hepatocarcinogen that, in combination with the choline-deficient diet, leads to hepatic fat loading, inflammation, fibrosis, hepatic progenitor cell (HPC) response, and eventually hepatocellular carcinoma development [5]. Proliferation is a key event in liver regeneration after injury. In a standard regeneration process, hepatocytes and cholangiocytes replenish their respective cell types and restore the hepatic mass. However, during chronic liver injury, when hepatocyte proliferation is inhibited or delayed, HPC aids in liver regeneration by first proliferating and later differentiating into functional hepatocytes [6]. Following CDE diet-induced injury, inflammation and activation of HPC are observed, making it a physiologically relevant model of chronic liver injury.

In this study, we investigated the role of HNF4α-cMyc interaction in CDE diet-induced liver injury. Our results demonstrated that deletion of HNF4α increases subacute liver injury induced by CDE diet feeding, and deletion of cMyc protects against the injury.

## Materials and Methods

### Animals, Treatment, and Tissue Harvesting

All animals were housed in facilities accredited by the Association for Assessment and Accreditation of Laboratory Animal Care at the University of Kansas Medical Center under a standard 12 h light/dark cycle with free access to chow and water. All studies were approved by the Institutional Animal Care and Use Committee at the University of Kansas Medical Center. The KUMC IACUC abides by the ARRIVE statement. Two-month-old male C57BL/6J mice were purchased from Jackson Laboratories and used for the initial analysis. HNF4α-floxed, cMyc-floxed and HNF4α-cMyc double floxed mice were injected intraperitoneally with AAV8-TBG-CRE to generate hepatocyte specific HNF4α-Knockout (KO), cMyc-KO and double-KO mice, respectively, as described previously [3]. Floxed mice treated with AAV8-TBG-GFP were used as controls. To study liver injury, these mice were given choline-deficient diet (Envigo) supplemented with 0.15% ethionine in drinking water (Acros Organics, 146170100) for one week. To study regeneration and recovery, mice were switched back to a normal chow diet and drinking water for one week after one week of a CDE diet. After euthanasia, liver and blood samples were collected and processed for further analysis, as described earlier [3].

### Protein isolation and western blot analysis

Western blot analysis was performed using whole liver lysates for HNF4α (Perseus Proteomics, PP-H1415-00) and cMyc (Cell Signaling Technology, 56055), as described previously [7]. GAPDH (Cell Signaling Technology, 2118) was used as a housekeeping control. Li-Cor Odyssey FC was used to image the Western blots.

### Real-Time Polymerase Chain Reaction

RNA isolation and its cDNA conversion were carried out as previously described [7]. qPCR analysis was done on BioRad CFX384. 100 ng of cDNA was used per reaction to measure the mRNA expression of genes of interest using the manufacturer’s protocol (ThermoFisher). The 18s gene expression in the same samples was used to normalize the ct values, as described previously [8]. The Ct values for the one-week CDE diet and recovery groups were compared to the normal diet given the control group of the respective phenotype. The primers used for real-time PCR are listed in Table 1.

### Staining Procedures

Paraffin embedded 5uM liver tissue sections were used for hematoxylin and eosin (H&E), immunohistochemical staining of HNF4α (Perseus Proteomics PP-H1415-00, 1:500), F4/80 (Cell signaling technology 70076, 1:200), αSMA (Cell signaling technology 19245, 1:500), Ki67 (Cell signaling technology 12202, 1:400), and immunofluorescence staining of Sox9 (Millipore Sigma AB5535, 1:1500) as previously described [7, 9]

### Serum ALT

Serum alanine aminotransferase (ALT) levels were measured using A Pointe Scientific ALT Assay kit by Fisher Scientific according to the manufacturer’s protocol.

### Statistical Analysis

Data presented in the form of bar graphs show the mean ± standard error of the mean. GraphPad Prism 9 was used to graph and calculate statistics. Two-way ANOVA and Student’s *t-test* were applied to all analyses, with *p*<0.05 considered significant. For all the experiments, number of 3-5 mice were used per group.

## Results

### Decreased HNF4α expression after one week of CDE diet

In the first experiment, two-month-old male C57BL/6J mice were fed a CDE diet for one week to study the injury response. To determine the regeneration response upon injury, these mice were allowed to recover on a normal diet for one week after the CDE diet (Fig. 1A). The CDE diet feeding resulted in an almost 10-fold increase in serum ALT levels, which were reduced after one week of recovery (Fig. 1B). Serum ALT data were corroborated by H&E staining, which showed disrupted liver histology with moderate fat accumulation and the presence of ductular reactions after CDE diet feeding. During the recovery period, mice showed restored histology similar to that of normal diet controls (Fig. 1C). *HNF4α* mRNA remained unchanged, but genes positively regulated by HNF4α including *Dio1* and *F12*, were significantly downregulated, whereas genes negatively regulated by HNF4α including *cMyc* and *Ect2*, were significantly upregulated after CDE diet feeding. During recovery on a normal diet. *HNF4α, Dio1* and *F12* mRNA expression returned to control levels, and *cMyc* and *Ect2* mRNA expression was reduced to those of normal diet controls (Fig. 1D-H). Consistent with these results, Western blot analysis showed decreased protein expression of HNF4α following the CDE diet (Fig. 1 I–J). The western blot results were validated by HNF4α immunohistochemistry (Fig. 1K).

**Figure 1:**
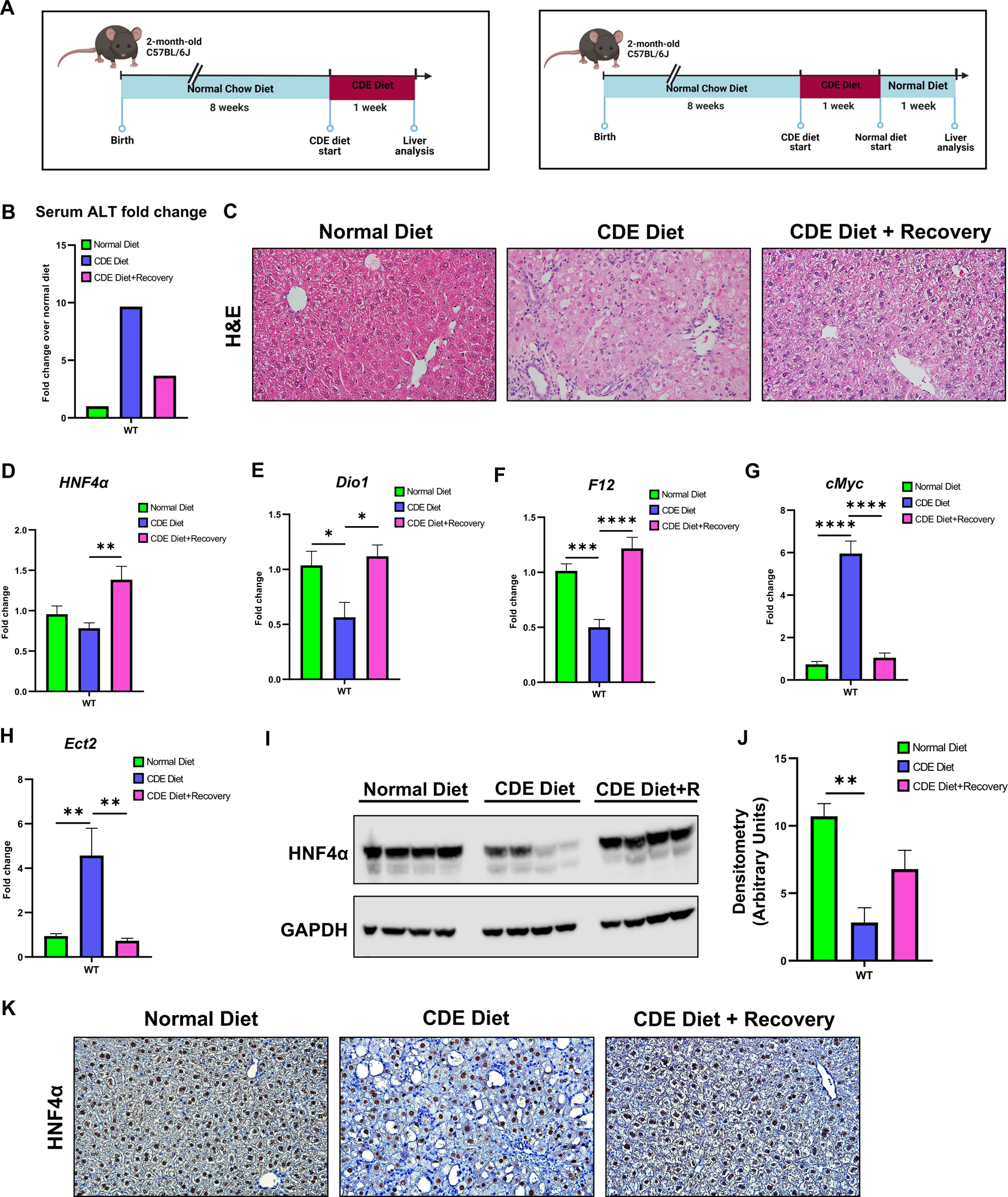
Decreased HNF4α expression after one-week of CDE diet. (A) Study design (B) Serum ALT fold change, (C) representative photomicrographs of H&E staining, real-time PCR analysis of (D) HNF4α, (E) Dio1, (F) F12, (G) cMyc and (H) Ect2, (I) Western blot analysis, (J) densitometric analysis for HNF4α and (K) representative photomicrographs of HNF4α immunohistochemistry of C57BL/6J mice after CDE diet induced injury and regeneration. Original magnification 400X. The bar represents the mean ± standard error of the mean. n=3-5. *p<0.05, **p<0.01, ***p<0.001, ****p<0.0001.

### Significantly higher liver injury following one week of CDE diet

Next, we studied liver injury and recovery in WT, HNF4α-KO, cMyc-KO and DKO mice after one week of CDE-diet feeding followed by one week of recovery on a normal diet (Fig. 2A-B). As previously published [10], HNF4α-KO mice showed an increased LW/BW ratio after CDE diet feeding, which was significantly reduced during the recovery phase (Fig. 2C). All four genotypes-WT, HNF4α-KO. The cMyc-KO and DKO mice showed significantly high serum ALT levels, indicative of increased liver injury, following one week of a CDE diet. HNF4α-KO mice showed higher injury, whereas the cMyc-KO mice experienced less injury than the WT mice. Injury was significantly resolved in all mice after one week of recovery on a normal diet (Fig. 2D). These findings were corroborated by H&E staining. Significant changes in histology, including fat accumulation observed in the staining images, were consistent with the injury data. The histological features were restored during the recovery phase (Fig. 2E).

**Figure 2:**
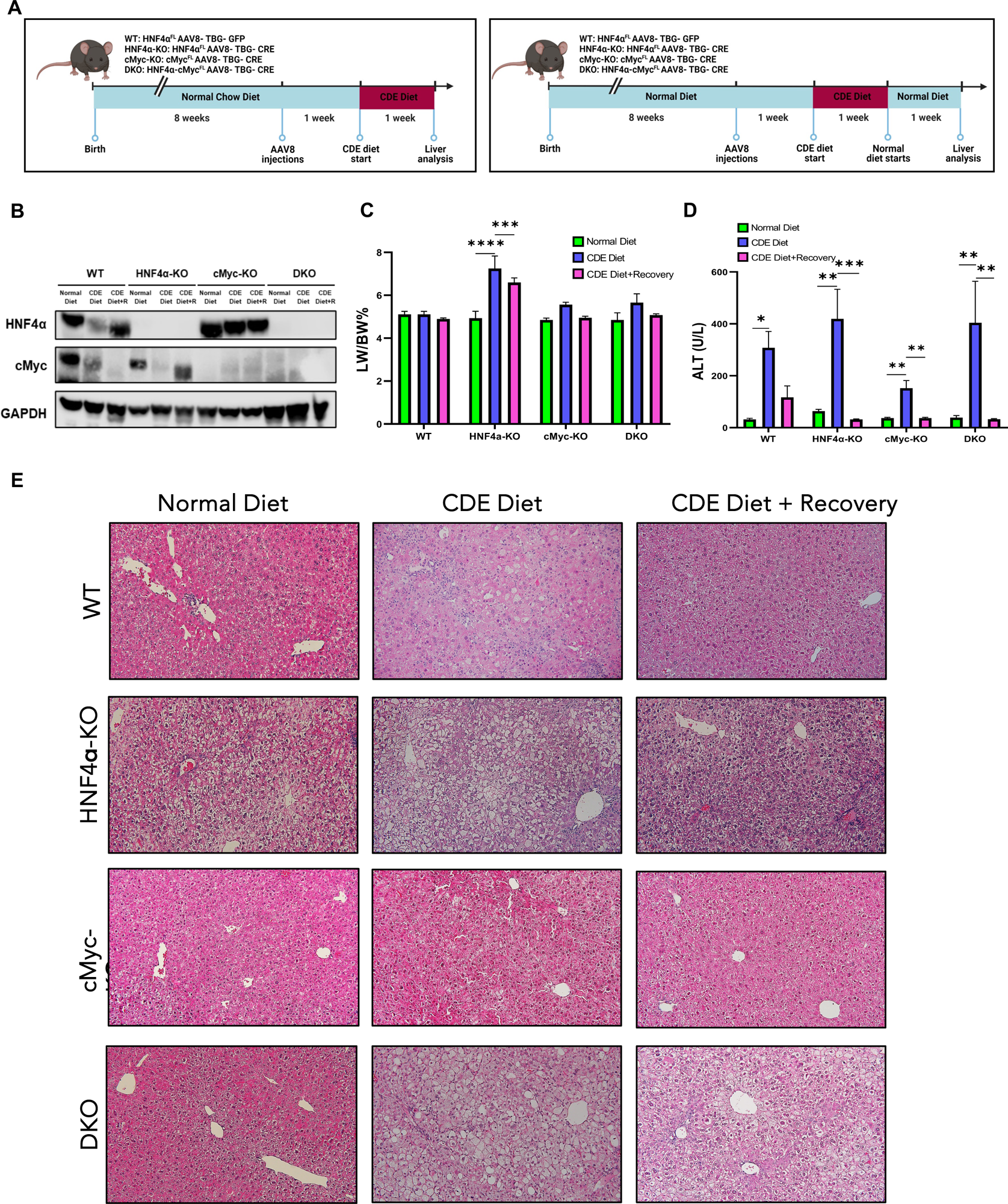
Significantly higher liver injury following one-week of CDE diet. (A) Study design (B) confirmatory Western blot analysis (C) Percent liver/body weight ratio, and (D) serum ALT levels, (E) representative photomicrographs of H&E staining in WT, HNF4α-KO, cMyc-KO and DKO mice after CDE diet induced injury and regeneration. Original magnification 200X. The bar represents the mean ± standard error of the mean. n=3-5. *p<0.05, **p<0.01, ***p<0.001, ****p<0.0001.

### Increased inflammation in cMyc-KO mice after one week of CDE diet

We determined the status of inflammation using qPCR analysis for several inflammation markers after one week of CDE diet feeding and one week of recovery. HNF4α-KO mice showed significantly higher expression of inflammation mediator genes, *including Il1b, Ifng, Tnfa,* and *Il10*, and gene markers of cells involved in inflammation response, including *Adgre1,Cd163, Clef4, Ccr2,* and *Vsig4,* during the injury phase. Expression of inflammatory genes was significantly higher in cMyc-KO mice compared to WT and HNF4α-KO mice. In DKO mice, the induction of inflammatory genes was comparable to HNF4α-KO mice (Fig. 3A). The mRNA data were further supported by F4/80 immunohistochemistry staining. Among the four genotypes, cMyc-KO mice showed significantly higher F4/80 positive cells after feeding the CDE diet (Fig. 3B). The increased mRNA and protein expression of inflammation markers during CDE diet-induced injury was significantly decreased during the recovery phase in all groups.

**Figure 3:**
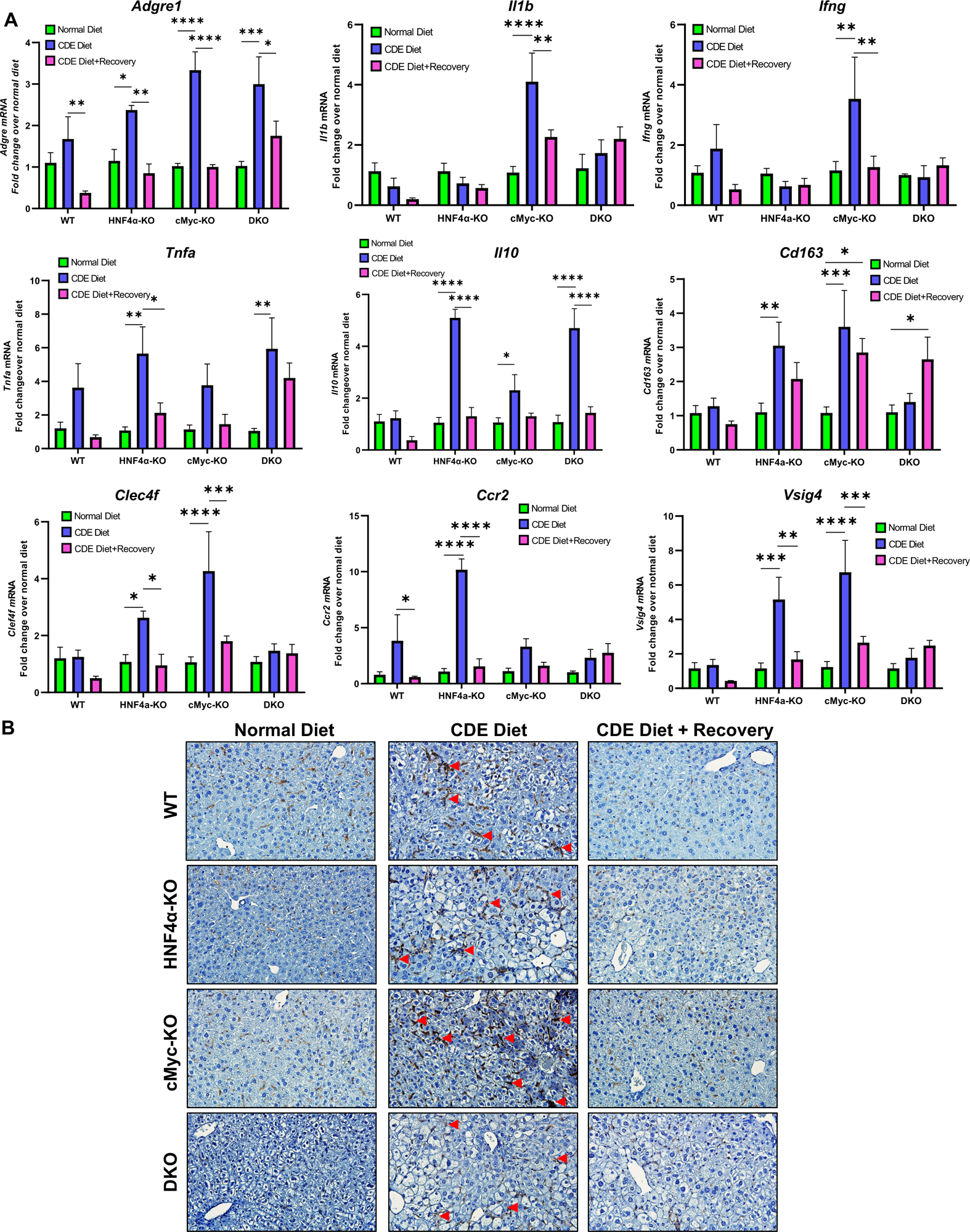
Increased inflammation in cMyc-KO mice after one-week of CDE diet. (A) Real-time PCR analysis of *Adgre*, *Il1b*, *Ifng*, *Tnfa, Il10, Cd163, Clef4f, Ccr2, Vsig4* and (B) representative photomicrographs of F4/80 immunohistochemistry in WT, HNF4α-KO, cMyc-KO and DKO mice after CDE diet induced injury and regeneration. Original magnification 400X. The bar represents the mean ± standard error of the mean. n=3-5. *p<0.05, **p<0.01, ***p<0.001, ****p<0.0001.

### Increased fibrosis in HNF4α-KO mice after one week of CDE diet

To investigate possible early fibrotic responses in these mice, we performed qPCR analysis of the known fibrosis markers, including *Des, Tgfb1*, *Col1a1*, *Col1a2,* and *Col3a1.* HNF4α-KO mice showed significantly higher expression of fibrosis genes during the injury phase. The cMyc-KO mice resulted in significant activation of collagen genes, which was lower than HNF4α-KO mice. In DKO, the induction of fibrosis genes was comparable to HNF4α-KO mice (Fig. 4A). The mRNA data were further corroborated by αSMA immunohistochemistry staining (Fig. 4B). During the recovery phase, increased fibrosis in HNF4α-KO and cMyc-KO mice significantly decreased. However, DKO mice showed consistently higher fibrosis responses, even during the recovery phase.

**Figure 4:**
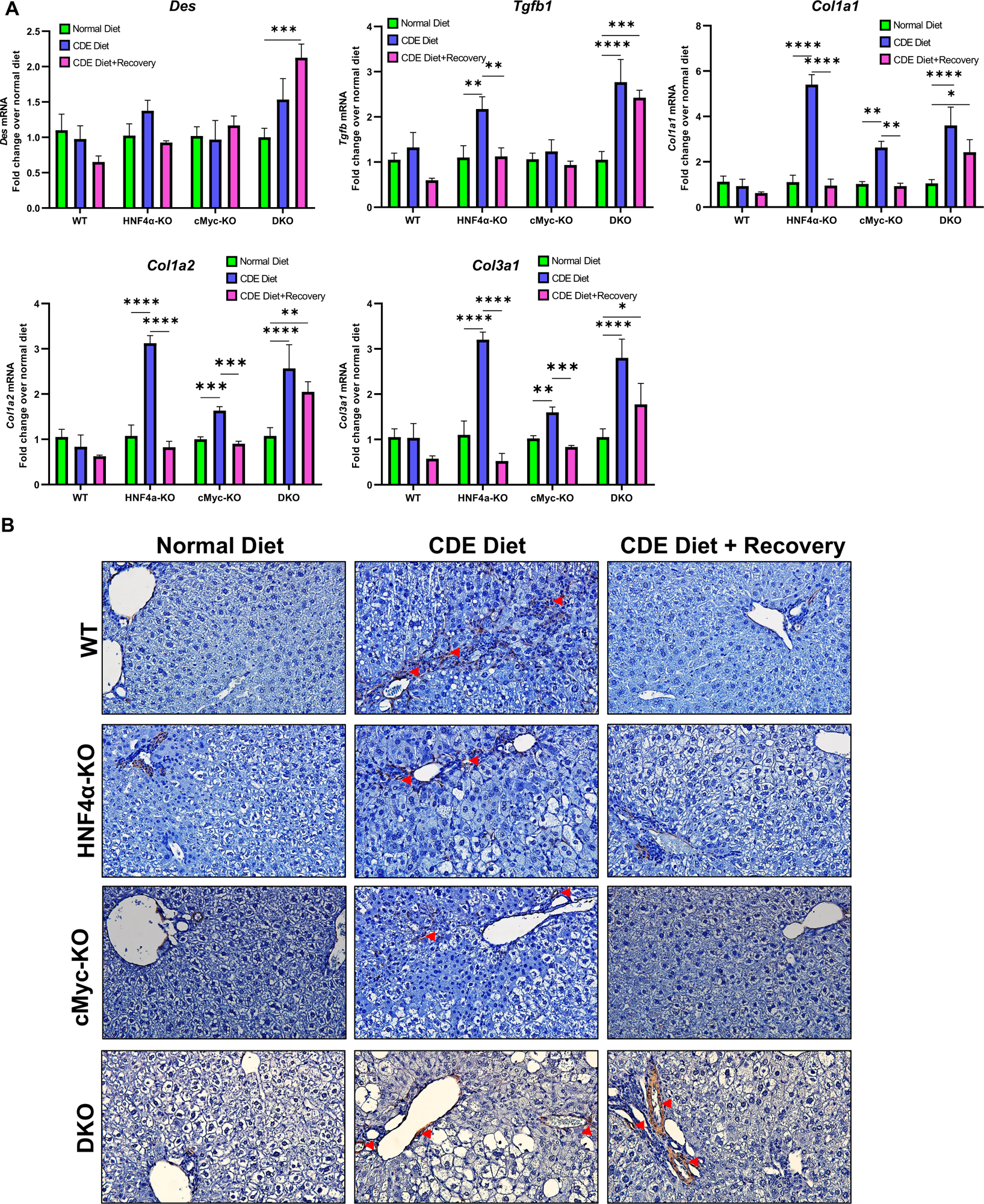
Increased fibrosis in HNF4α-KO mice after one-week of CDE diet. (A) Real-time PCR analysis of fibrosis markers *Des, Tgfb*, *Col1a1*, *Col1a2,* and *Col3a1* and (B) representative photomicrographs of αSMA immunohistochemistry in WT, HNF4α-KO, cMyc-KO and DKO mice after CDE diet induced injury and regeneration. Original magnification 400X. The bar represents the mean ± standard error of the mean. n=3-5. *p<0.05, **p<0.01, ***p<0.001, ****p<0.0001.

### Increased proliferation in HNF4α-KO mice after one week of CDE diet

We investigated liver regeneration following CDE-diet-induced liver injury by qPCR analysis of proliferation markers including *Ccnb1, Ccnd1,* and *Ccne1.* WT mice exhibited minimal changes in proliferation genes, but HNF4α-KO, cMyc-KO and DKO mice showed a significantly higher expression of Cyclins during the injury phase. The increase in cyclin expression was reduced back to control levels in cMyc-KO mice during recovery. However, HNF4α-KO and DKO mice showed continued expression of cyclin genes, albeit lower than during the injury phase (Fig. 5A). Ki67 immunohistochemistry further confirmed an increased proliferation response (Fig. 5B).

**Figure 5:**
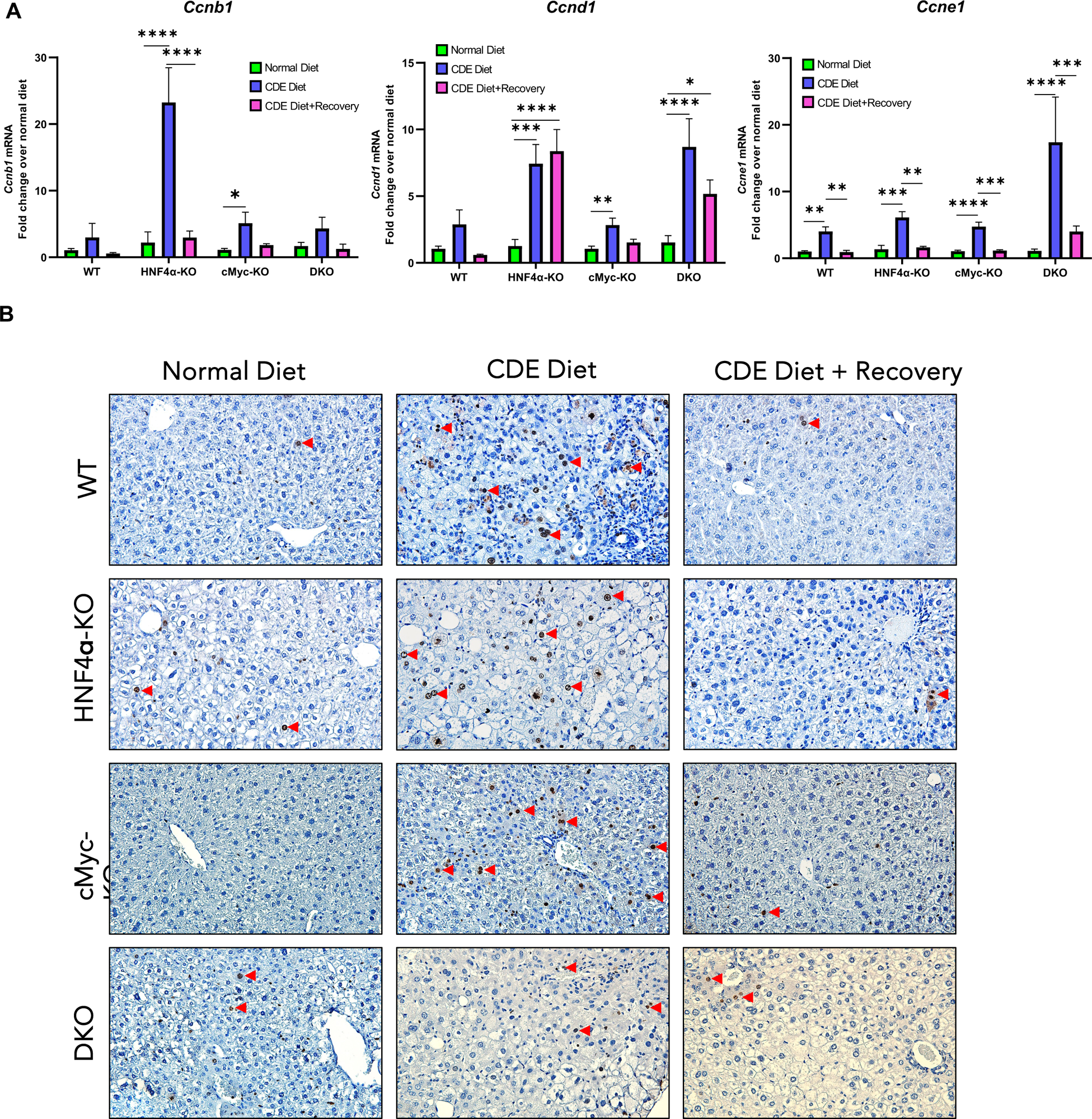
Increased proliferation in HNF4α-KO mice after one-week of CDE diet. (A) Real-time PCR analysis of proliferation markers *Ccnb1, Ccnd1* and *Ccne1* and (B) representative photomicrographs of Ki67 immunohistochemistry in WT, HNF4α-KO, cMyc-KO and DKO mice after CDE diet induced injury and regeneration. Original magnification 400X. The bar represents the mean ± standard error of the mean. n=3-5. *p<0.05, **p<0.01, ***p<0.001, ****p<0.0001.

### Significant increase in HPC markers both following one week of CDE-induced injury and after one week of recovery in DKO mice

Finally, to investigate the HPC activation and HPC-driven proliferation response in these mice, qPCR analysis of HPC markers, such as *Krt19*, *Epcam,* and *Sox9,* was performed. WT mice exhibited no change in *Krt19* or *Epcam* but showed an increase in *Sox9* expression only during the injury phase. HNF4α-KO mice showed an increase in *Krt19* and *Epcam* expression only during the injury phase. cMyc-KO mice showed no change in *Epcam* and *Sox9* expression, but a small increase in Krt19 only during the recovery phase. The DKO mice showed the most robust HPC activation response with an increase in *Krt19* and *Epcam* expression during the recovery phase and an increase in Sox9 during both the injury and recovery phases (Fig. 6A). Sox9 immunofluorescence further supported the mRNA data, showing increased Sox9 positive cells in all groups following a CDE diet for one week (Fig. 6B). The DKO mice had the highest number of Sox9-positive cells in the liver after one week of CDE diet feeding. Interestingly, we did not detect any Sox9-positive cells during the recovery phase in DKO livers despite significantly higher *Sox9* mRNA expression.

**Figure 6:**
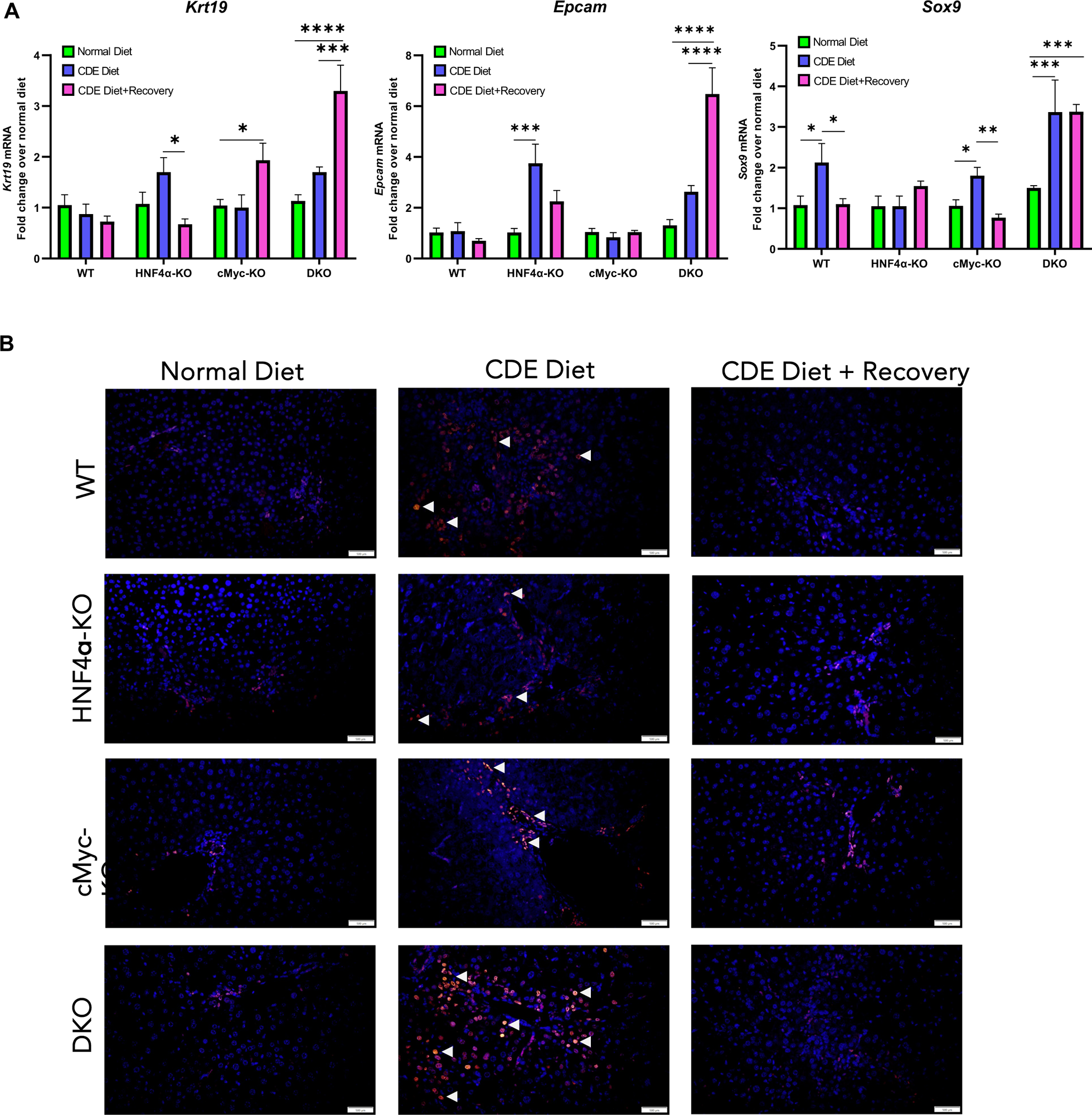
Significant increase in HPC markers both following one week of CDE-induced injury and after one week of recovery in DKO mice. (A) Real-time PCR analysis of *Krt19*, *Epcam* and *Sox9*, and (B) representative photomicrographs of SOX9 immunofluorescence in WT, HNF4α-KO, cMyc-KO and DKO mice after CDE diet induced injury and regeneration. Original magnification 400X. The bar represents the mean ± standard error of the mean. n= 3-5. *p<0.05, **p<0.01, ***p<0.001, ****p<0.0001.

## Discussion

The role of HNF4α, an orphan nuclear receptor expressed exclusively in hepatocytes, has been well-defined in embryonic liver development, as well as for post-natal maintenance of the differentiated phenotype [2]. Recent studies have shown that HNF4α also inhibits hepatocyte proliferation and maintains a differentiated state. Loss of HNF4α results in de-differentiation and increased proliferation in the absence of liver injury [2]. Studies from our laboratory and others have shown that progression of chronic liver disease progression is associated with loss of HNF4α activity [11]. Recent studies from our laboratory explored the role of HNF4α in liver regeneration after partial hepatectomy and after acetaminophen overdose, a model of drug-induced liver injury [3, 4]. The partial hepatectomy model has relatively low injury, no inflammation, and synchronized liver regeneration, while the APAP overdose model has necrosis and inflammation followed by unsynchronized liver regeneration. However, in both models, injury is induced acutely. Further, these models do not have either the early fibrotic changes or activation of HPCs, which are the hallmark of chronic liver disease. The goal of this study was to investigate the role of HNF4α in liver regeneration, subacute liver injury, and subsequent regeneration after CDE diet feeding in which inflammation, fibrosis, and HPC-mediated proliferation are involved.

In our initial experiment with 2-month-old C57BL/6J mice, mice experienced 10 times higher liver injury as compared to the control mice following one week of a CDE diet. We observed a significant decrease in HNF4α expression at the mRNA and protein levels in these mice. However, these mice showed a significant reduction in liver injury and re-gained HNF4α expression upon recovery on a normal diet. These data suggest that maintaining HNF4α expression is necessary for recovery from CDE-diet-induced liver injury. CDE diet studies in hepatocyte-specific HNF4α-KO mice supported these findings. HNF4α-KO mice experienced higher liver injury after CDE diet feeding compared to WT mice. These mice, however, recovered and restored their histology after one week of recovery. Quantification of inflammation and fibrosis markers revealed that HNF4α-KO mice had higher expression of these genes during injury compared to WT mice. This finding further highlighted the significance of maintaining HNF4α expression during subacute liver injury. To analyze the regeneration process in these mice, we studied the proliferation response. Deletion of HNF4α is known to induce a spontaneous proliferation response in hepatocytes [10]. Consistent with these findings, HNF4α-KO mice showed significant induction in the expression of cyclins and Ki67-positive cells during the injury period. These data suggest that decreased HNF4α expression during injury is associated with increased hepatocyte entry into the cell cycle, helping these mice recover via regeneration.

Deletion of HNF4α results in increased expression of cMyc at the mRNA and protein levels [2, 3]. cMyc is a proto-oncogene that is often deregulated in various cancers [12]. cMyc is a major regulator of cell proliferation, cell cycle progression, cell growth, metabolism, and differentiation [13]. The spontaneous proliferation and metabolic changes observed following HNF4α deletion are at least in part due to upregulation of cMyc. Therefore, we investigated the role of HNF4α-cMyc interaction in liver regeneration after subacute liver injury induced by CDE diet feeding. Deletion of both HNF4α and cMyc in DKO mice resulted in significant liver injury comparable to the HNF4α-KO mice after one week of CDE diet feeding and complete recovery on a normal diet for one week. Increased inflammation, fibrosis, and proliferation response after CDE diet feeding were higher than in WT mice and comparable to HNF4α-KO mice. DKO mice showed consistently higher fibrosis responses during the recovery phase. Interestingly, DKO mice showed a significant increase in HPC markers both following one week of CDE-induced injury and after one week of recovery. This suggests that in the case of both HNF4α and cMyc deletion, liver regeneration and recovery are driven less by hepatocyte proliferation and more by HPC activation. These data support our hypothesis that cMyc activation after HNF4α induces hepatocytes proliferation.

Our recent studies showed that the deletion of cMyc in HNF4α-KO mice reduced acetaminophen-induced acute liver injury [4]. However, in the current study of CDE diet-induced injury, where the injury is physiologically similar to that of chronic liver diseases, we found that deletion of cMyc in HNF4α-KO mice exacerbated HPC-driven proliferation. In humans, increased HPC activation is associated with the severity of chronic liver disease [14]. This further supports the hypothesis that during chronic liver injury, when HNF4α expression is low, cMyc might play a compensatory role in aiding recovery.

In summary, deletion of HNF4α increases subacute liver injury induced by CDE diet feeding, and deletion of cMyc protects against the injury. Deletion of HNF4α results in increased inflammation, fibrosis, proliferation, and HPC activation, all of which except inflammation are reduced following cMyc deletion. Simultaneous deletion of HNF4α and cMyc results in a phenotype similar to HNF4α deletion, however, involves HPC-driven regeneration response to recover from CDE diet-induced injury. Taken together, these data show that HNF4α protects against inflammatory and fibrotic change following CDE diet-induced injury, which is driven by cMyc.

---

**Supplementary Table 1:**
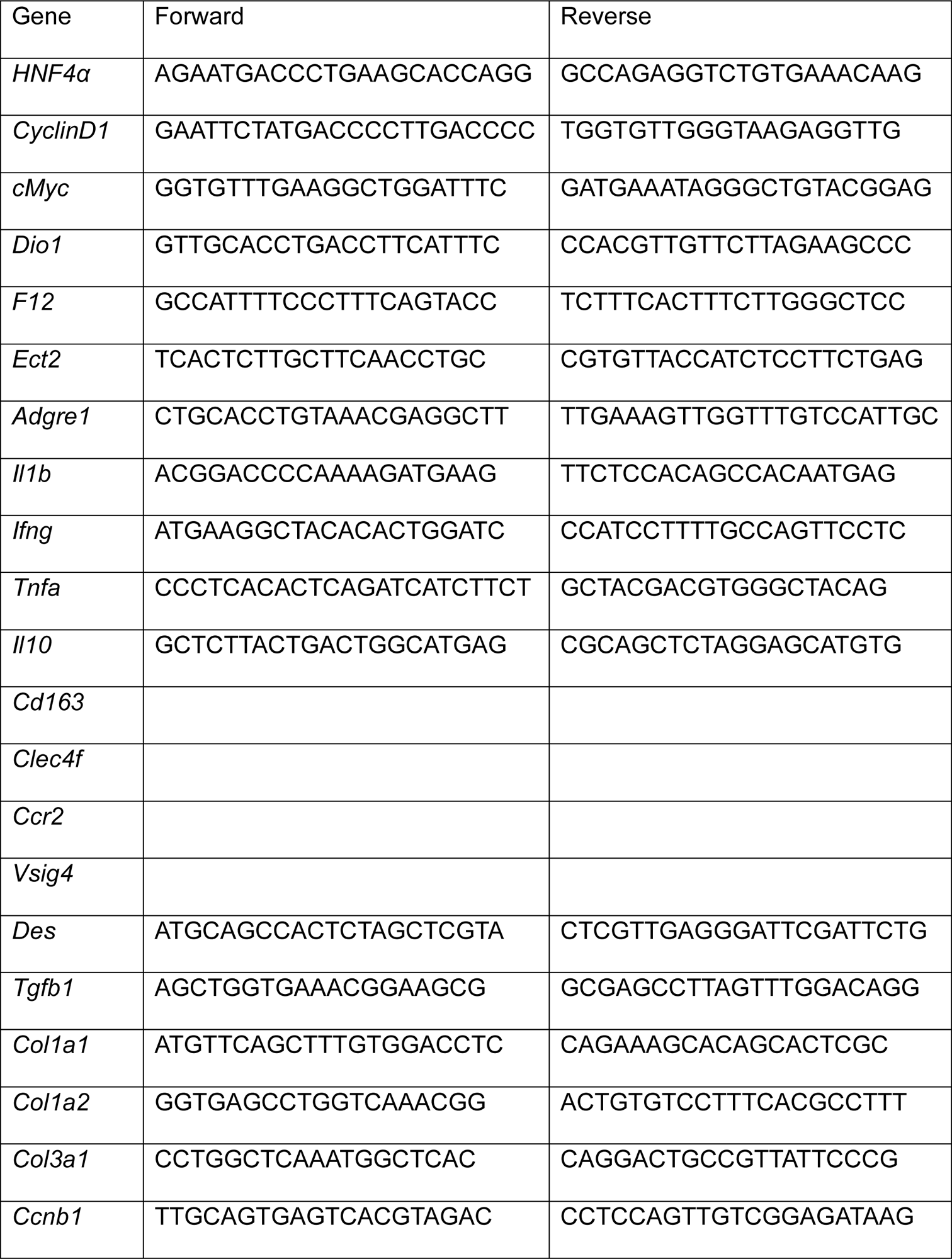
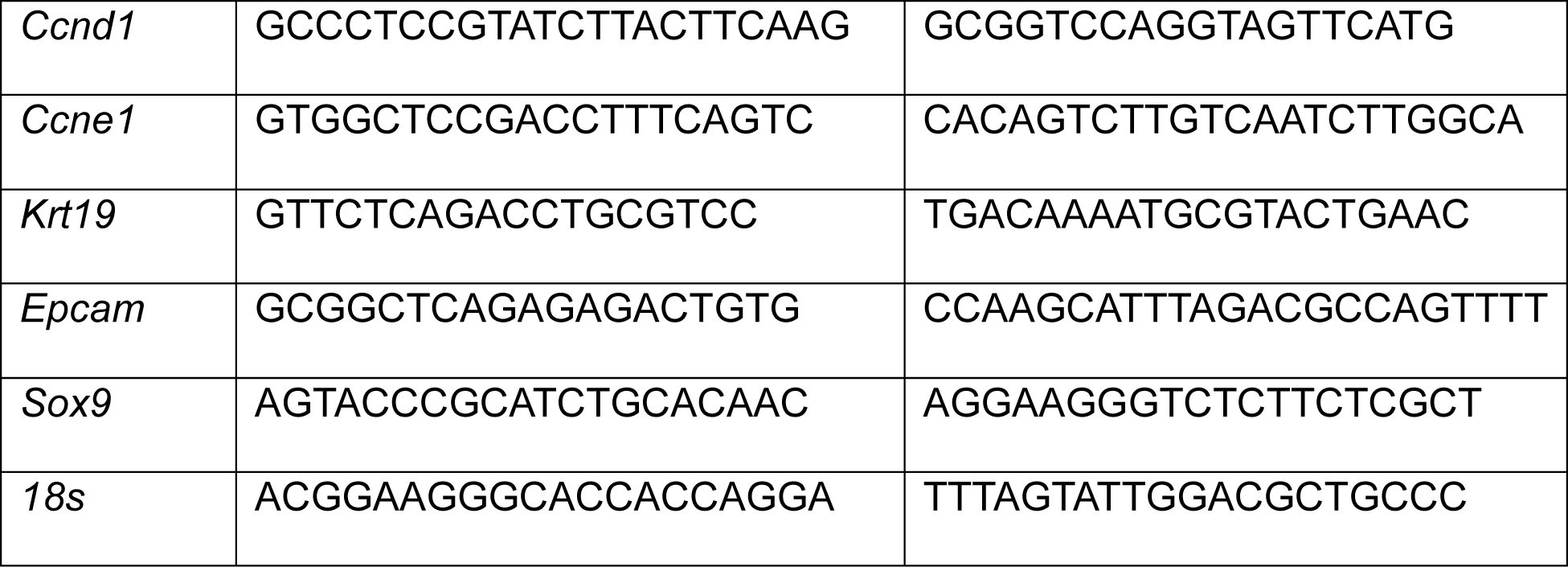
Primers used in this study.

